# Regulation of protein abundance in neurons by selective translation of 3′UTR isoforms

**DOI:** 10.64898/2026.07.08.737200

**Authors:** Sakshi Gorey, Hasan Can Ozbulut, Judit Carrasco, Ying Zhang, Anton Hess, Ipek Akol, Carlos Alfonso-Gonzalez, Mengjin Shi, Dominika Grzejda, Julian Wolter-Mess, Mirijam Egg, Fernando Mateos, Sarah Holec, Alejandro Gomez-Auli, Nina Cabezas-Wallscheid, Tanja Vogel, Sabine Rospert, Valérie Hilgers

## Abstract

The precise regulation of protein synthesis is essential for cellular function and survival. In particular, in neurons, dysregulated mRNA translation is linked to impaired memory formation and is a hallmark of neurodegenerative diseases. Neurons are characterized by tissue-specific, long 3′ untranslated regions (3′UTRs); in this study, we demonstrate that mRNA isoforms with these neuronal 3′UTRs are less efficiently translated than their short counterparts in Drosophila and mammalian brains. 3′UTR-dependent translation is based on a negative feedback mechanism centered around the two neural-enriched proteins ELAV and Pumilio. The long *elav* 3′UTR inhibits production of the neuronal ELAV protein, which in turn mediates 3′UTR extension of hundreds of neuronal genes. Those long 3′UTR isoforms are preferentially bound and translationally inhibited by Pumilio. The regulatory loop maintains optimal neuronal 3′UTR and protein levels in conditions of genetic and environmental perturbations; its disruption reduces animal viability and lowers stress resilience, and causes severe developmental phenotypes in flies and in human brain organoids. We propose 3′UTR-mediated translational control as an evolutionarily conserved mechanism for the maintenance of cell-type-specific proteostasis.

## Introduction

Neurons are long-lived cells whose functionality strongly depends on the precise regulation of protein abundance and localization, and in which disruption of proteostasis is particularly detrimental, causing neurodegeneration^1,2^. Spatiotemporal control of protein synthesis is a major mechanism for specifying proteomes in brain development, and transcripts are increasingly translationally regulated as neurons form and mature^3,4^. Accordingly, specific proteome identities of nervous system cell types are shaped primarily through translational regulation of a defined subset of transcripts^5,6^. Neuronal proteomes also differ by subcompartment and even synapse subtype, at least in part due to local and spatial modulation of protein synthesis^7,8^. This uncoupling of mRNA availability and protein abundance enables neurons to respond to signaling cues and adapt to genetic and environmental fluctuations. Disruption of translation control in neural tissues is associated with neurodevelopmental and neurodegenerative disorders, and inhibiting *de novo* protein synthesis in specific cell types of the brain impairs long-term threat memory consolidation^9–11^.

The association of mRNAs with translating ribosomes is predominantly regulated by RNA-binding proteins (RBPs) that bind to sequence elements in 3′ untranslated regions (3′UTRs)^12^. mRNA variants of the same gene that differ by their 3′UTR (3′UTR isoforms) can also differ in their translational output^13,14^, and recent studies have identified genes whose 3′UTR isoforms vary in their translation rates in separate cell types of the mouse brain^15^. The differential translation of 3′UTR isoforms of individual genes has been implicated in oncogenic activation^16,17^ and neuronal signaling^18,19^, showing that ratios between short and long 3′UTR isoforms help define cellular protein output.

Neurons possess a particularly extensive 3′UTR diversity due to cell-type specific RBPs that regulate alternative polyadenylation (APA) during neurogenesis. In neural stem/progenitor cells, PQBP1 promotes the preferential use of proximal polyadenylation sites (PASs) of cell cycle genes, thereby maintaining short 3′UTR profiles in proliferative cells^20^. Conversely, in neurons, ELAV/Hu proteins bind to and inhibit proximal PASs in pre-mRNAs of hundreds of genes, promoting cleavage at more distal sites and the generation of extended 3′UTRs^21–24^. The emergence of mRNAs with longer, neuron-specific 3′UTR sequences, thereafter referred to as nUTRs, has been documented in various animals from flies to mammals including humans^25–33^. Through their additional binding sites for RBPs and microRNAs, nUTRs have the potential to confer differential properties to distinct 3′UTR isoforms of the same gene. Generally, mRNAs with longer 3′UTRs are more stable, more localized, and less translated^6,13,34^. Genes with several isoforms in neurons tend to encode proteins associated with specialized neuronal functions such as synaptic transmission; from those genes, mRNA isoforms with longer 3′UTRs tend to be more localized^34^. The ratio of 3′UTR isoform synthesis can be significantly altered by enhanced activity^34^. These studies raise the possibility that the modulation of 3′UTR extension through APA helps neurons to maintain proteostasis and to adapt protein levels in response to intrinsic and extrinsic fluctuations.

Here, we interrogate the broader role of nUTRs in the regulation of protein output in the nervous system. We demonstrate that in fly heads, mouse brains and human neural progenitor cells, mRNAs with longer 3′UTRs are generally less translated compared to shorter isoforms of the same gene. This differential translation efficiency is crucial to maintain physiological levels of neuronal proteins, and ensures survival in conditions of genetic and environmental fluctuations. Mechanistically, nUTR-mediated proteostasis is regulated through two processes: i) an autoregulatory feedback loop in which the nUTR of the neuronal gene *elav* keeps ELAV protein at precise levels and, consequently, preserves the nUTR balance of hundreds of neuronal genes, ii) the nUTR-specific translational repression of those genes by the RBP Pumilio. Our work demonstrates the important role and mechanism of the selective translation of 3′UTR isoforms in cell-type-specific protein abundance.

## Results

### Long 3′UTR isoforms of neuronal genes are less translated than their short counterparts

We investigated the isoform-specific translation of nervous system mRNAs by polysome profiling, which we performed in mouse brains^35^ and adapted for fly heads (see methods). The method separates cytoplasmic components based on their sedimentation in a sucrose gradient, yielding fractions containing RNAs with increasing numbers of bound ribosomes. RNA-seq on each fraction enabled the identification of translating and non-translating mRNAs at the isoform level (Fig. 1A). In a principal component analysis (PCA) of gene expression, fractions clearly clustered by biological replicates and according to their translational status (Fig. S1A, B). As a control, we investigated mRNAs encoding the “housekeeping” proteins Gapdh and Actin, which were progressively enriched with increasing ribosome density in both Drosophila and mouse brains (Fig. S1C, D), consistent with an effective translation of cell-essential genes. To compare the translation state of distinct 3′UTR isoforms of the same gene, we determined, for each fraction, the abundance of long compared to short 3′UTR isoforms using QAPA^36^ (Table S1). We found that 30% and over 50% of all expressed APA genes (i.e., genes expressing at least two APA isoforms) in flies and mice, respectively, display a bias in which either the long or the short 3′UTR isoform was enriched in translating fractions compared to lysate (Fig. 1B). These genes exhibited a clear pattern: in the actively translating fractions, isoform expression was shifted towards shorter 3′UTRs, and long 3′UTR isoforms were overrepresented in the non-translating fraction (Fig. 1C, D).

**Figure 1.**
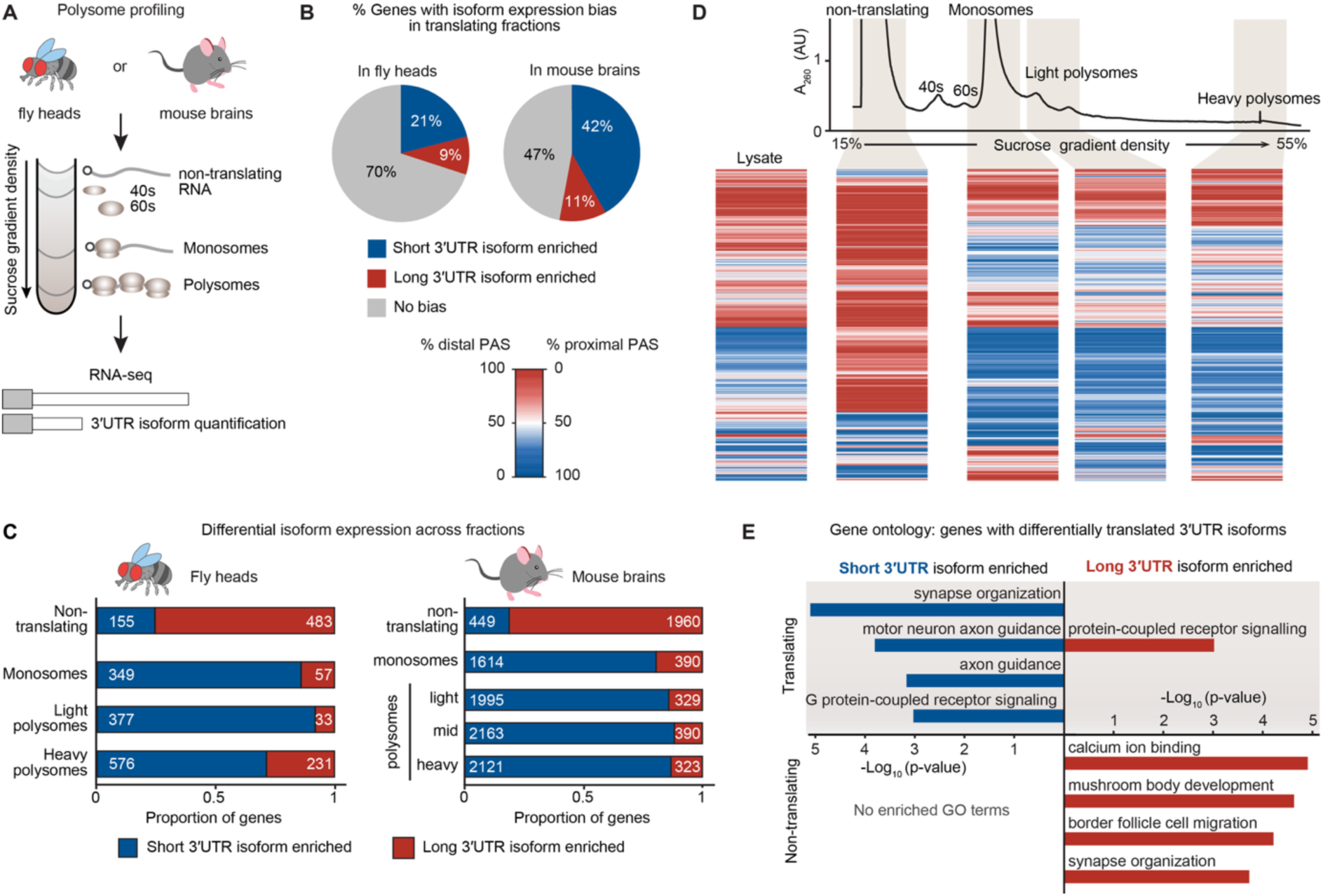
Long 3′UTR isoforms of neuronal genes are less translated than their short counterparts. **(A)** Workflow of polysome profiling performed in adult Drosophila heads and mouse brains. A lysate prepared from freshly separated fly heads or mouse brains in the presence of cycloheximide was subjected to ultracentrifugation in a sucrose gradient, in which the number of ribosomes bound to RNAs increases with density. RNA was isolated from each fraction for total RNA sequencing to calculate the translation status of individual mRNA isoforms, based on the expression of each transcript relative to lysate. **(B)** Differential translation of short or long 3’UTR isoforms for each gene in fly heads and mouse brains. Shown are genes in which the indicated 3’UTR isoforms (short or long) are enriched in the translating fractions. **(C)** Proportion of genes in fly heads and mouse brains that display differential translation of 3’UTR isoforms. Values indicate the number of genes for which isoform expression ratios are biased (compared to lysate) towards the short (blue) or long (red) 3’UTR isoform. **(D)** Heat maps representing the differential usage of proximal or distal poly(A) sites in the indicated polysome profile fractions, in 1064 Drosophila genes clustered by Ward’s method. All genes with differential translation of 3’UTR isoforms are shown, with one line representing one gene across all fractions. Relative amounts of short and long 3’UTRs are represented by relative poly(A) site usage, with 50% signifying equal relative abundance of both isoforms. A_260_, UV absorbance of the gradient at 260nm (arbitrary unit). **(E)** Gene ontology analysis of genes for which short (blue) or long (red) 3’UTR isoforms are enriched in either translating or non-translating fractions from fly heads. The top four terms are shown for each category. Background set: all expressed genes.

A gene ontology (GO) analysis revealed significant enrichment of GO terms for short isoforms in the translating fraction and long isoforms in non-translating fractions: neuron development and synaptic signal transduction. In contrast, few significant terms were found for highly translated long isoforms or non-translating short isoforms (Fig. 1E, Fig. S1E–F, Table S2). This could indicate that processes related to neuronal differentiation and synapse functionality are regulated through 3′UTR-isoform-dependent translational repression.

### The *elav* neuronal 3′UTR regulates translation and ELAV protein abundance in neurons

We tested in flies whether long 3′UTRs reduce translation efficiency of the mRNA that carries them, using the model gene *elav* (Fig. 2A). We chose *elav* because; i) its encoded protein ELAV promotes nUTR production in hundreds of genes, placing it at the hierarchical summit of 3′UTR regulation in neurons; ii) *elav* and its human orthologue *ELAVL1* are themselves regulated at the 3′UTR level^37,38^, which may indicate a conserved mechanism of ELAV autoregulation. In our polysome profile from fly heads, *elav* behaved similarly to most other multi-UTR genes: the long *elav* 3′UTR isoform was translated substantially less efficiently compared to the short (Fig. 2B). This result was surprising because ELAV protein expression is exclusive to cells that express the long, neuron-specific *elav* 3′UTR^21^, and studies have shown mechanisms of miRNA-mediated repression of the short *elav* 3′UTR in non-neural cells^39^. To investigate whether longer 3′UTRs and lower translation are causally linked, we deleted the approximately 3 kb-long extended portion of the *elav* 3′UTR (the nUTR) using CRISPR-Cas9-mediated gene editing (Fig. 2A). In the resulting *elav^ΔnUTR^* flies, ELAV protein was mildly but consistently and significantly upregulated without any increase in *elav* RNA levels (Fig. 2C, Fig. S2A, B), showing that the *elav* mRNA can be efficiently translated in absence of the nUTR.

**Figure 2.**
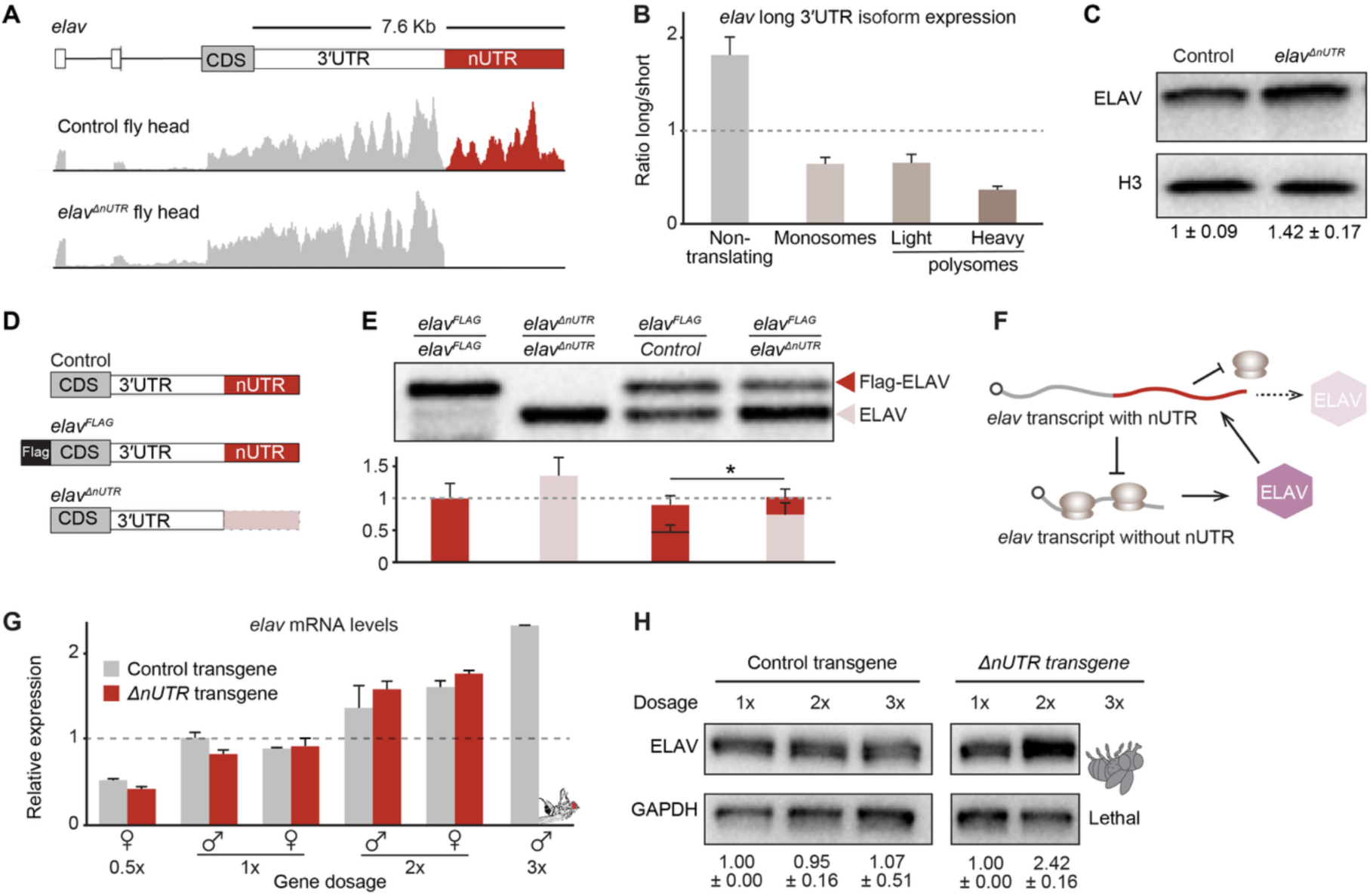
The neuronal 3’UTR regulates mRNA translation and maintains optimal ELAV protein levels. **(A)** *elav* gene model (drawn to scale) and total RNA-seq tracks in fly heads of control and *elav^ΔnUTR^* flies. *elav^ΔnUTR^* flies are homozygous for an *elav* allele lacking the neuronal 3’UTR. **(B)** RT-qPCR quantification of the expression of the long *elav* 3’UTR isoform relative to total (short) *elav* mRNA in fly head polysome profile fractions, normalized to levels in tissue lysate. Error bars represent mean ±SD of two biological replicates. **(C)** ELAV protein expression in control and *elav^ΔnUTR^* adult fly heads. ELAV band intensity was normalized to the respective Histone H3 band intensity, with ELAV levels in control flies set to 1. Mean ±SD is indicated for three biological replicates. Ten heads of 3–5-day-old flies were used per replicate. **(D)** *elav* alleles used to create the fly genotypes used in (**E**). In *elav^FLAG^*, a Flag tag was inserted into the endogenous *elav* locus to generate a visibly larger protein. **(E)** ELAV protein expression by Western blot (top) and relative band intensity quantification (bottom) in adult fly heads of the indicated genotypes. Each genotype (all females) consists of two of the alleles shown in **(D)**. The proportion of ELAV expressed from each allele (control and *elav^FLAG^*, red; or *elav^ΔnUTR^*, light pink) was normalized to total ELAV per lane, with ELAV levels in *elav^FLAG^/elav^FLAG^*set to 1. Error bars represent mean ±SD of three biological replicates. *p ≤ 0.05 (Welch’s t-test). **(F)** Model of 3’UTR-mediated regulation of *elav* translation, in which mRNA isoforms containing the longer 3’UTR (nUTR, red) are translationally repressed and short isoforms produce the bulk of the encoded protein. ELAV protein in turn promotes the formation of longer 3’UTRs, negatively feeding back on ELAV abundance. **(G, H)** *elav* RNA (**G**, males and females) and ELAV protein (**H**, males) expression in adult heads of flies carrying increasing numbers of *elav* transgenes, to obtain the indicated gene dosage. Transgenes consisted of either the wild-type *elav* gene region (control) or lacked the *elav* nUTR (*ΔnUTR*). Genotypes are as follows, with + denoting a wild-type chromosome and *tsg* a chromosome carrying a transgene. Males, 1x (+/Y ; ; +/+), 2x (*+/Y ; ; tsg/*+), 3x (*+/Y ; ; tsg/tsg*). Females, 0.5x (*Δelav/+ ; ; +/+)*, 1x (+/+ ; ; +/+), 2x (*+/+; ; tsg/tsg*). For RT-qPCR, *elav* mRNA levels were normalized to *RpL32* mRNA and levels in flies “control transgene, 1x dosage” were set to 1. For Western Blot, ELAV band intensity was normalized to the respective GAPDH band intensity, with ELAV levels in flies “1x dosage” set to 1. Error bars represent mean ±SD of three biological replicates for each genotype. Ten heads of adult flies were used per replicate.

We next directly compared protein produced from the endogenous *elav* locus with and without nUTR, by combining, in the same fly, the *elav^ΔnUTR^* allele with an endogenously Flag-tagged wild-type *elav* allele, which produces a distinguishably larger Flag-ELAV protein (Fig. 2D). ELAV production from *elav*^FLAG^ was equal to that of an untagged, wild-type allele, in a fly expressing these two control alleles. In contrast, when expressed with *elav*^FLAG^, *elav^ΔnUTR^* predominantly contributed to ELAV protein production (Fig. 2E). This notable visible expression difference between the two alleles still likely overrepresents the contribution of the long 3′UTR isoform, because the wild-type allele produces both short and long isoforms (Fig. 2A). Together with our previous results, this experiment demonstrates that the long, nUTR-containing *elav* isoform is translationally repressed. The short isoform efficiently produces ELAV, whose molecular function consists in promoting 3′UTR extension, including that of its own nUTR. Therefore, here we describe a negative feedback loop in which ELAV promotes the formation of translationally repressed *elav* mRNA isoforms, which in turn reduces ELAV levels (Fig. 2F).

### A highly effective feedback loop preventing detrimental ELAV overexpression

To challenge the functionality and buffering capacity of this regulatory loop, we created conditions of strong genetic fluctuations *in vivo*. We duplicated a 100 kb genomic region of the X chromosome encompassing the *elav* transcription unit and introduced it onto the third chromosome. We created an *elav^ΔnUTR^* version of this transgene by removing the *elav* nUTR by BAC recombineering (as in Fig. 2D). Since the *elav* genomic region is subject to dosage compensation by approximately two-fold overexpression in males (who carry only one X chromosome), we could vary *elav* gene dosage from 0.5-fold to two-fold in female flies, and from one-fold to three-fold in male flies (see details in Fig. 2G legend). Increases in *elav* gene dosage caused proportional increases of *elav* mRNA levels, in a manner similar for wild-type and *elav^ΔnUTR^* transgenes (Fig. 2G). However, for protein levels, effects of *elav* overexpression were nUTR-dependent: with wild-type transgenes, ELAV levels remained constant even when gene dosage was increased three-fold, showing that ELAV protein expression is tightly regulated, and can withstand substantial genetic variation. In contrast, with *elav^ΔnUTR^* transgenes, ELAV was overexpressed, and a three-fold dosage increase caused 100% lethality (Fig. 2H).

We attempted to shift the *elav* 3′UTR isoform balance towards overexpression of the long form, through several gene editing approaches in flies. First, we deleted the key proximal *elav* PASs, abrogating cleavage at those sites, to force transcriptional read-through and distal PAS use; second, we introduced a self-cleaving hammerhead ribozyme sequence^40^ designed to specifically deplete long *elav* transcripts (Fig. S2C). All mutations substantially reduced expression of the long, nUTR-carrying *elav* transcripts in fly brains; but this reduction was insufficient to counteract *elav* autoregulation, and ELAV protein levels were indistinguishable from those of control flies (Fig. S2D, E). Multiple fail-safe mechanisms ensure proper *elav* transcription and processing^37,41^, likely supporting a robust feedback loop. Overall, our findings reveal that 3′UTR-dependent regulation is efficient at maintaining constant protein levels even in a strongly fluctuating genetic environment and can prevent lethal protein overexpression.

### 3′UTR-dependent translational regulation maintains proteostasis during development

To assess whether 3′UTR-dependent translation control is not only efficient at buffering strong fluctuations, but also fine-tunes protein levels relevant in physiological contexts, we characterized *elav^ΔnUTR^* flies, in which ELAV is only mildly overexpressed (by 30-40%, Fig. S2B). *elav^ΔnUTR^* flies display reduced overall viability (Fig. 3A) and suffer developmental delays (Fig. 3B, C). These defects, though not evident without quantification, are likely highly detrimental to fitness and survival outside of laboratory conditions.

**Figure 3.**
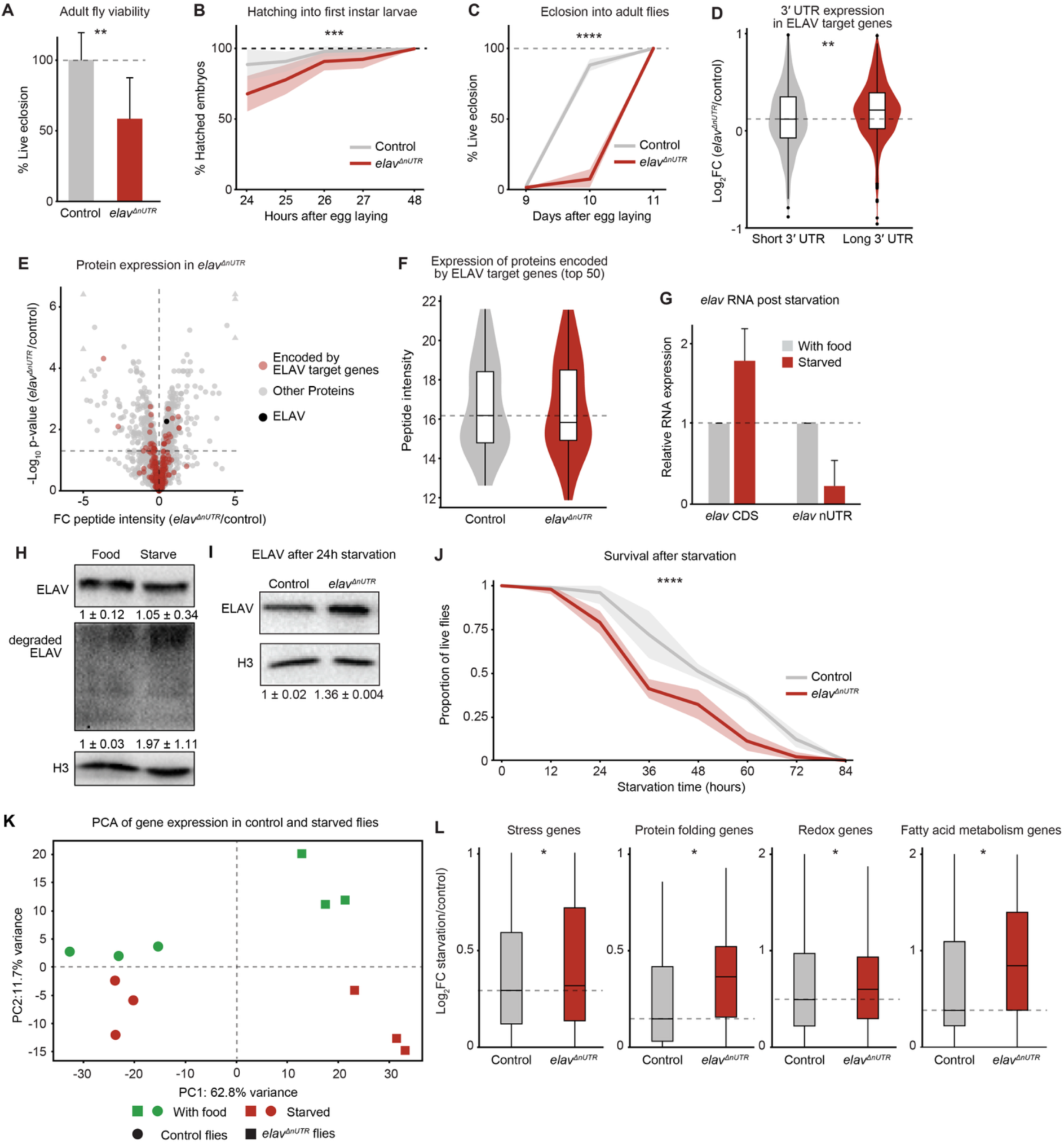
3′UTR-dependent translational regulation maintains proteostasis during development and stress. **(A)** Viability of *elav^ΔnUTR^* flies, measured as the rate of eclosion of live adult flies, compared to expected Mendelian ratios. Eclosion rates were normalized to those of control flies. Error bars indicate mean ±SD of four biological replicates. 375 (control) and 414 (*elav^ΔnUTR^*) adult flies were scored. **p ≤ 0.01 (Welch’s t-test). **(B, C)** Fraction of *elav^ΔnUTR^* and control flies that hatched from embryos into first instar larvae **(B)**, and that eclosed from pupae into live adults **(C)**, at the indicated time points after egg laying. Mean ±SEM of 10 biological replicates (10 embryos/pupae per replicate) is represented for each genotype. χ²=12.93, p=3.23 e-0.4 **(B)**, χ²=133.9, p=<0.0001 **(C)** (Mantel-Cox log-rank test). **(D)** Differential expression of the short (500 bp preceding the pPAS) and long 3′UTR in 313 ELAV target genes (genes that undergo ELAV-mediated 3′UTR extension), in *elav^ΔnUTR^* compared to control fly heads. Expression was quantified from RNA-seq data. ** p≤0.01 (Wilcoxon test). **(E)** Differential protein expression in *elav^ΔnUTR^* compared to control fly heads. Peptide intensity was measured by mass spectrometry. Proteins encoded by ELAV target genes are marked in red, with ELAV in black. **(F)** Expression of proteins encoded by ELAV target genes (top 50 genes with long 3’UTR isoform upregulated), in *elav^ΔnUTR^* compared to control fly heads. **(G)** RT-qPCR quantification of the long and short *elav* 3’UTR isoforms in adult heads of flies kept in control (with food) conditions or under starvation stress for 24h (starved). RNA levels were normalized to *RpL32* mRNA and levels in control conditions were set to 1. Error bars represent mean ±SD of three biological replicates. **(H, I)** ELAV protein expression in adult heads of flies that were subjected to starvation stress for 24h. In **(H),** the Western blot membrane was cut below the ELAV band at 50 kDa and exposed separately to visualize degradation products. Intensities of the ELAV band and the ELAV degradation smear were normalized to the respective Histone H3 band intensity, with levels in control conditions set to 1. Mean ±SD is indicated for four **(H)** and three **(I)** biological replicates. **(J)** Stress resilience of control and *elav^ΔnUTR^*flies, measured as the proportion of flies alive after being subjected to starvation stress for the indicated number of hours. Mean ±SEM of five biological replicates (10 males and 10 females per replicate) is represented for each genotype. χ²=23.12, p=1.5 X10^−6^ (Mantel-Cox log-rank test). **(K)** Principal component analysis of gene expression in adult heads of control and *elav^ΔnUTR^* flies kept in control (with food) conditions or under starvation stress for 24h (starved). **(L)** Differential expression of genes of the indicated gene sets in adult heads of flies kept under starvation stress for 24h compared to control fly heads. Expression was quantified from RNA-seq data. * p≤0.05 (Wilcoxon test).

ELAV inhibits proximal PAS usage in neurons, strongly shifting 3′UTR isoform ratios towards long, nUTR-containing mRNAs in hundreds of genes, here referred to as “ELAV APA targets”^21,22^. We tested whether a moderate ELAV overexpression has an impact on cellular nUTRs. In heads of *elav^ΔnUTR^* flies, 3′UTRs of ELAV APA targets were globally —but mildly—lengthened (Fig. 3D, Fig. S3A), indicating that the loss of the *elav* nUTR increases not only ELAV levels, but also its cellular activity. We measured protein abundance in control and *elav^ΔnUTR^* fly heads by mass spectrometry (Table S3). Consistent with long 3′UTRs negatively regulating translation of neuronal proteins, levels of proteins encoded by these genes were downregulated in *elav^ΔnUTR^* flies, although the difference was not statistically significant (Fig. 3E, F).

### Loss of neuronal 3′UTR balance increases vulnerability to starvation stress

Regulating protein abundance is essential during stress, particularly in the brain, and several mechanisms safeguard the proteome under stress^42^. In organisms from plants to mammals, alteration of mRNA 3′UTR length has been observed in response to dehydration^43^, starvation, arsenic^44^ and ribotoxic^45^ stress, and APA has been proposed to function as an adaptive mechanism to preserve mRNAs in response to stress. Together with our results, this points to the interesting possibility that nUTR-dependent regulation of translation could contribute to protein homeostasis and stress resilience in the animal nervous system. We used intermittent food shortage —a natural challenge in an animal’s life— as a physiological condition to test this hypothesis. After 24h of starvation, we observed a drastic shortening of the *elav* 3′UTR in the brain of adult flies (Fig. 3G). This was accompanied by substantial degradation of ELAV protein. Levels of intact ELAV however, remained indistinguishable from normal conditions (Fig. 3H). Independently of which event occurred first —protein degradation or 3′UTR shortening, both hallmarks of stress— maintenance of cellular ELAV abundance could be explained by an increase in ELAV production stemming from the upregulation of the translationally active, short *elav* 3′UTR isoform, thereby compensating for ELAV degradation. Importantly, in contrast to that of *elav*, other 3′UTRs remained relatively stable, and *elav* APA target genes displayed moderate or no change in 3′UTR length (Fig. S3B, C), consistent with levels of functionally active ELAV being maintained under starvation. *elav^ΔnUTR^*flies, in which 3′UTR-mediated translation regulation is impaired, and ELAV is upregulated (Fig. 3I), died significantly faster than controls under conditions of starvation (Fig. 3J). Upon stress exposure, stress-related genes were more upregulated in *elav^ΔnUTR^*flies compared to control flies (Fig. 3K, L, Fig. S3D), indicating that dysregulation of the cellular nUTR levels impairs the stress response. In conclusion, our data show that 3′UTR mediated translational regulation constitutes an important mechanism for proteostasis and supports survival during physiological challenges.

### Mechanism of nUTR-dependent translational repression

Neuron-enriched, longer 3′UTRs often carry conserved binding motifs for Pumilio (Pum)^25,26,46^, a highly conserved RBP known to repress translation, with roles in synaptic function across species^47–50^. Pum was recently shown to regulate the expression of synaptic genes by binding to, and retaining, long 3′UTR isoforms in the neuronal soma in fly and mammalian neurons^46,51^. In light of the preferential translation of short 3′UTR isoforms of neuronal genes (Fig. 1), we hypothesize that Pum specifically binds and translationally represses long, nUTR-carrying mRNAs. Using flies in which Pum was endogenously Flag-tagged, we performed RNA immunoprecipitation followed by RNA-seq (xRIP-seq) using a Flag antibody; Pum binding specificity was verified by comparison with lack of signal in control (untagged) flies. We found extensive Pum binding signal over the entire *elav* 3′UTR (Fig. 4A, Fig. S4A), indicating the RBP may translationally repress both long and short *elav* mRNA isoforms. To test this, we assessed ELAV expression in brains of *pumilio* mutant flies (*Δpum*). Since loss of Pum is lethal, we used a combination of hypomorphic alleles viable until the third instar larval stage. In *Δpum* brains, ELAV was strongly upregulated relative to control brains (Fig. 4B). Interestingly, even though Pum binding did not detectably differ between *elav* long and short isoforms (Fig. 4A), long 3′UTR *elav* isoforms were significantly more sensitive to Pum loss than short ones: ELAV expression from the nUTR-containing wild-type *elav* allele, but not the *elav^ΔnUTR^* allele, was drastically elevated in *Δpum* mutants (Fig. 4C), showing that Pum regulates ELAV protein output in an 3′UTR-specific manner.

**Figure 4.**
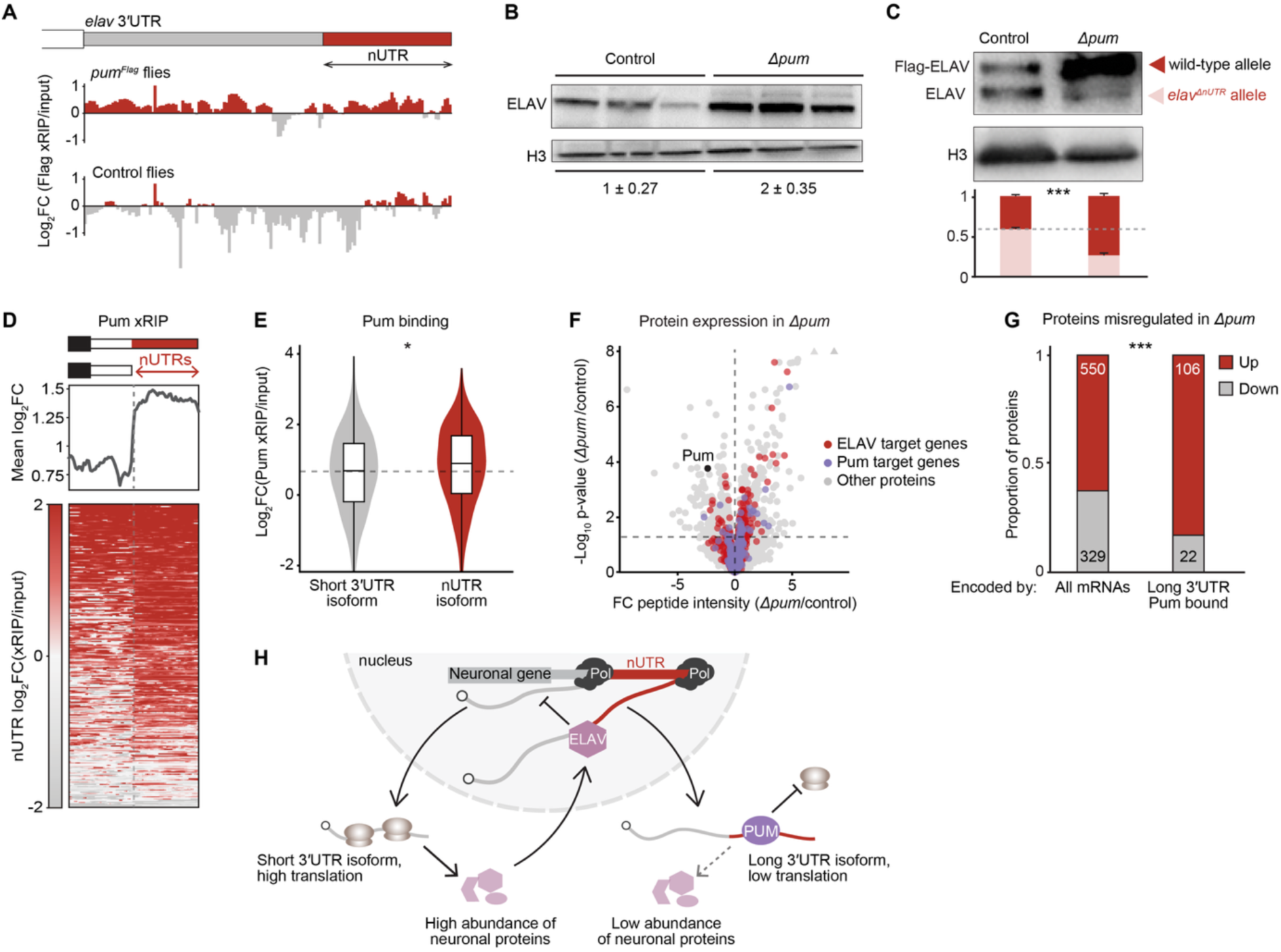
Mechanism of nUTR-dependent translational repression. **(A)** Flag xRIP-seq signal tracks for the *elav* 3′UTR, in flies in which Pumilio was endogenously Flag-tagged (*pum^Flag^*), and in untagged *control* flies *(w^1118^)*. Signal is shown normalized to respective input. **(B)** ELAV protein expression in brains of control and *Δpum* larvae. Flies are homozygous for the *elav^FLAG^* allele on the first chromosome. **(C)** Western blot (top) and relative band intensity quantification (bottom) of ELAV protein in brains of control and *Δpum* larvae. All larvae are females expressing both Flag-ELAV (from the *elav^FLAG^* allele, red arrowhead) and untagged ELAV (from the *elav^ΔnUTR^* allele, light pink arrowhead). Histone H3 serves as a loading control. The proportion of ELAV expressed from each allele was normalized to total ELAV per lane, with ELAV levels in Control set to 1. Error bars represent mean ±SD of three biological replicates. *** p ≤ 0.001 (Welch’s t-test). **(D)** Enrichment of neuronal 3′UTRs in Pum xRIP relative to input. The heatmap profile plot displays 1 kb upstream, and the neuronal 3′UTR downstream (scaled region, indicated in red), of the proximal poly(A) site. Pum xRIP was performed in adult fly heads. **(E)** Quantification of Pum binding to short and nUTR-containing mRNA isoforms of ELAV target genes, by Pum xRIP-seq signal compared to input. p=0.0438 (Wilcoxon test). **(F)** Differential protein expression in *Δpum* compared to control fly heads. Peptide intensity was measured by mass spectrometry. Proteins encoded by Pum target genes (mRNAs enriched in Pum xRIP-seq compared to input) and proteins encoded by ELAV target genes (genes that undergo ELAV-mediated 3′UTR extension) are marked in purple and red, respectively. Pum is black. **(G)** Proportion and numbers of proteins encoded by transcripts of the indicated groups that are upregulated (up) and downregulated (down) in *Δpum* mutant larval brains compared to control. All mRNAs are compared to mRNAs whose long 3’UTR isoform is preferentially bound by Pum compared to the short isoform of the same gene. **(H)** Model of nUTR-mediated translational regulation of neuronal genes. nUTR-containing mRNAs are translationally repressed. Low levels of ELAV protein promotes expression of short 3’UTR isoforms, which in turn increases global protein abundance. The feedback loop is supported by nUTR-specific binding and translational repression by Pumilio.

At the global level, performing an isoform-aware quantification of Pum xRIP-seq signal, we found that Pum displays a strong binding preference for long, translationally repressed 3′UTR isoforms of neuronal genes (Fig. 4D, E, Fig. S4B, C), raising the interesting hypothesis that Pum regulates the protein abundance of neuronal genes by repressing the translation of long 3′UTR isoforms. We quantified protein levels in *Δpum* larval brains by mass spectrometry. We found 550 proteins upregulated and 329 down, 23% and 5% of which encoded by Pum mRNA targets, respectively, indicating that Pum translationally represses the transcripts it binds in larval brains (Table S4, Fig. 4F). At the isoform level, neuronal genes in which 3′UTR isoforms were differentially bound by Pum were highly enriched among proteins upregulated in *Δpum* brains (Fig. 4G). Therefore, our data provide strong evidence that Pum translationally represses long 3′UTR isoforms of neuronal genes.

Taken together, we reveal a novel mode of protein output regulation, through a top-down mechanism, with ELAV dictating 3′UTR length of neuronal mRNAs and Pum regulating translation in an nUTR-dependent manner (Fig. 4H).

### Neurodevelopmental impairment in human brain organoids lacking the *ELAVL1* nUTR

Mouse and human neurons express four homologues of the ELAV/Hu protein family, ELAVL1–4 (also known as HuA or HuR, HuB, HuC, and HuD). The precise spatiotemporal expression of ELAVL proteins, crucial for proper neuronal development, is tightly regulated at the transcriptional and post-transcriptional level; dysregulation is associated with neurodevelopmental disorders^52–54^. Human *ELAVL1* mRNAs, like Drosophila *elav*, exist in the form of short and long 3′UTR isoforms, with the long 3′UTR enriched in neurons. In transgene experiments in human cells, *ELAVL1* APA and 3′UTR extension was mediated by ELAVL proteins^38,55^. Based on this, we speculate a role for the *ELAVL1* nUTR in neuronal protein homeostasis and mammalian brain development.

We used human induced pluripotent stem cells (iPSCs) and differentiated them into neural stem cells (NSCs) and neurons. Similar to Drosophila *elav*, expression of the long *ELAVL1* 3′UTR isoform gradually increases in the course of differentiation from iPSC to NSC and is most pronounced in neurons (Fig. 5A). We performed polysome fractionation in NSCs, and found that, for most genes, short 3′UTR isoforms are more efficiently translated compared to their long counterpart (Fig. 5B, C, Fig. S5A), following the global trend for neuronal genes in fly heads and mouse brains (Fig. 1). Accordingly, by qPCR, the *ELAVL1* nUTR isoform was underrepresented in translating NSC fractions (Fig. 5D). This result is consistent with studies showing that an *ELAVL1* transgene carrying the short 3′UTR produces more protein than the full-length version^38,55^. To assess the role of the *ELAVL1* nUTR in an *in vitro* model of human brain development, we deleted 2.1 kb of it using CRISPR-Cas9 gene editing (Fig. 5A), obtaining two independent *ELAVL1^ΔnUTR^* mutant human iPSC cell lines. We differentiated the mutants and control iPSCs into cerebral organoids for 34 days (Fig. S5B, C).

**Figure 5.**
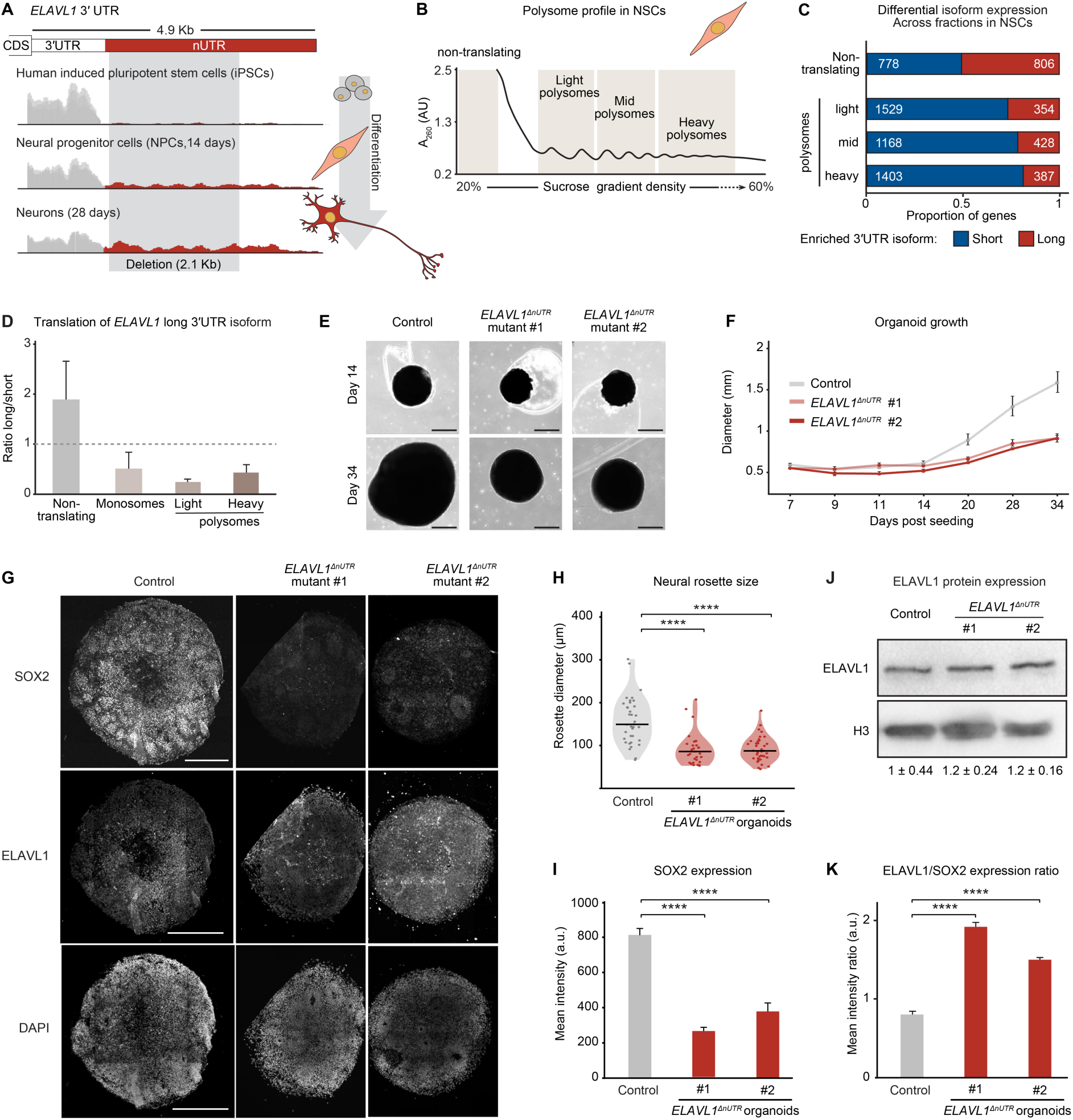
Neurodevelopmental impairment in human brain organoids lacking the ELAVL1 nUTR. **(A)** Gene model and mRNA-seq tracks of the *ELAVL1* 3’UTR (drawn to scale) in human iPSCs, and in the course of differentiation into neural progenitor cells and neurons. **(B)** Profile from NSC polysome profiling experiment. The four fractions used for downstream RNA-seq analysis are indicated. A_260_, UV absorbance of the gradient at 260nm (arbitrary unit). **(C)** Proportion of genes in NSCs that display differential translation of 3’UTR isoforms. Values indicate the number of genes for which isoform expression ratios are biased (compared to lysate) towards the short (blue) or long (red) 3’UTR isoform. **(D)** RT-qPCR quantification of the expression of the long *ELAVL1* 3’UTR isoform relative to total (short) *elav* mRNA isoforms in polysome profile fractions, normalized to levels in tissue lysate. Error bars represent mean ±SD of three biological replicates. **(E)** Light microscopy images of cerebral organoids grown from control and *ELAVL1^ΔnUTR^* human iPSCs on day 14 and day 34. Scale bar: 500µm. **(F)** Mean diameter of control and *ELAVL1^ΔnUTR^* (two independent mutants) cerebral organoids at the indicated number of days post-seeding. Error bars represent mean ±SD for six organoids per genotype and time point. **(G)** Apotome imaging of control and *ELAVL1^ΔnUTR^*cerebral organoids stained for SOX2, ELAVL1 and DAPI. Scale bar: 400µm. **(H)** Quantification of neural rosette size in control and *ELAVL1^ΔnUTR^* cerebral organoids. 36 (control), 31 (mutant #1) and 40 (mutant #2) rosettes were scored. ****p < 0.0001 (Mann-Whitney U test). **(I)** Quantification of the mean signal intensity for SOX2 (background intensity-corrected) in control and *ELAVL1^ΔnUTR^* organoids. Error bars represent mean ±SD for four replicates. ****p < 0.0001 (Mann-Whitney U test). **(J)** Western blot comparing ELAVL1 protein expression in control and *ELAVL1^ΔnUTR^* day 5 organoids. ELAVL1 band intensities were normalized to the respective Histone H3 band intensity, with ELAVL1 levels in control organoids set to 1. Mean ±SD is indicated for three biological replicates. **(K)** Quantification of the ELAVL1/SOX2 signal intensity ratio in control and *ELAVL1^ΔnUTR^* organoids. Error bars represent mean ±SD for four replicates. *p < 0.05 (Welch’s t-test).

Organoids lacking the *ELAVL1* nUTR displayed severe developmental phenotypes. They appeared to lose structural integrity by day 11 (Fig. S5D) and failed to grow normally (Fig. 5E, F, Fig. S5B). In addition, *ELAVL1^ΔnUTR^*organoids displayed significantly smaller rosettes (Fig. 5G, H) and lower expression of the pluripotency marker SOX2 (Fig. 5G, I), suggesting that mutant organoids undergo premature differentiation. ELAVL1 was expressed broadly across cell types; notably, in *ELAVL1^ΔnUTR^* organoids, ELAVL1 was upregulated in SOX2-negative cells (Fig. 5J, K), indicating that the nUTR mediates *ELAVL1* translational repression specifically in more differentiated neural cells. It is likely that *ELAVL1* derepression in organoids lacking the *ELAVL1* nUTR biases cells towards more neuronal fates, causing severe growth and differentiation impairments.

While we provide strong evidence for nUTR-dependent autoregulation of ELAV/Hu proteins in flies and in a human model of early neural development, we cannot exclude the possibility that in the mammalian system, phenotypes are not directly due to translational regulation of nUTR-containing transcripts. Moreover, ELAVL proteins including ELAVL1 were shown to directly regulate translation in the cytoplasm^56^, implicating a more complex network regulating the abundance of neuronal proteins.

## Discussion

In this study, we establish APA as a global, conserved, gene-specific and tissue-specific regulator of translational efficiency and proteostasis. Most long 3′UTRs function as repressive, tunable modules that limit protein output. An autoregulatory loop involving the mutual inhibition between the long and short *elav* mRNA isoforms dictates 3′UTR length and translation efficiency of hundreds of genes, thereby maintaining neuronal proteins at precise physiological levels.

Deficiencies in ELAV protein levels *in vivo* are very robustly rescued by multiple mechanisms^41^, including stop-codon read-through^57^, exon-activated rescue by the paralogue *fne*^22^, functional redundancies with other ELAV/Hu homologues^23,58^ and the nUTR-dependent translational modulation reported here. Consequently, fly models with drastically reduced ELAV levels are fully viable, with neurodevelopment phenotypes of variable severity, and mild ELAV downregulation causes no detectable defects^59^. In contrast, the mild ELAV overexpression achieved here by deleting the *elav* nUTR causes viability loss, developmental delays and lack of resilience in stress conditions. This reveals a surprising cellular vulnerability towards gene overexpression, and may imply that nUTRs represent the central, perhaps only global instrument safeguarding neural tissues against excessive neuronal protein abundance. The protective role played by long 3′UTRs in constraining translation under specific physiological conditions likely extends beyond the extreme APA environment of the nervous system. Long-standing research shows extensive APA dysregulation in many diseases including but not limited to neurological disorders. For example, 3’UTR shortening is causally associated with oncogenic protein production in cancer cells, through mechanisms that are not entirely clear^12,16,60^.

The repressive role of the *elav* nUTR on ELAV protein production was unexpected: intuitively, the well-described neuron-restricted expression of both the *elav* nUTR and ELAV protein points to the nUTR-containing isoform as translated in neurons, especially since the short *elav* 3′UTR is expressed in other cells of the nervous system, such as neuroblasts and glia—where ELAV levels are very low^21^. Work from the Lai lab showed that both long and short *elav* 3′UTR isoforms are repressed by microRNAs and the RBP Mei-P26 outside of neurons^39^, a repression that could include non-neural cells of the nervous system. Together with this, our work identifying distinct functions for the two *elav* isoforms in neurons reconciles the apparent discrepancies between mRNA and protein expression patterns, and highlights the impact of gene 3′UTRs and post-transcriptional regulation mechanisms on determining when and where a protein is expressed.

Our finding that nUTR-mediated translational repression is mediated by Pumilio complements recent models for spatiotemporal tuning of neuronal proteins. In fly^46^ and mouse^51^ neurons, long 3′UTR isoforms of synaptic genes are selectively retained in neuronal cell bodies by Pum, while short 3′UTR isoforms are transported to the synapse, presumably for translation. Together with our study, this indicates that nUTRs play an important role in modulating localized translation of synaptic genes. The coordination of distinct post-transcriptional mechanisms through a common set of sequences and RBPs may help coordinate expression output in a specialized cellular environment. In light of findings that transcripts of functionally related groups of genes, for example transcription factors, are often co-localized in subcellular, membraneless compartments^61^, we postulate that neuronal 3′UTRs evolved to co-regulate the localization and translation efficiency of genes that execute highly specialized neuronal functions.

It will be interesting to identify how long neuronal 3’UTRs integrate information of RBP binding, RNA structure, condensation, and RNA modifications to mediate function^8^. Work on long 3′UTRs in neurons found a positive correlation between mRNA stability, translation, and localization, with more stable 3′UTRs, which are often longer, more likely to diffuse over long distances to form a distinct localized pattern^62,63^. These processes involve a myriad of other RBPs, with longer 3′UTRs expanding the number and possible combinations of RBP binding sites^64^. Our work shows that nUTRs are co-regulated by ELAV and Pum in the nervous system; of note, other APA genes, including those displaying differential isoform translation, are not Pum targets, indicating that Pum uniquely regulates neuronal processes. Some of Pum’s known roles in axonal transport, neuronal excitability, and synaptic growth^49,65–67^, could be due to this differential regulation, since many nUTR-bearing genes encode proteins involved in these processes. Hence, the question remains which RBPs and other factors regulate APA isoforms in other genes, and whether modules of tissue-specific co-regulation exist.

Through nUTR-dependent translational regulation, and other isoform-specific mechanisms, the expression of distinct mRNA isoforms may guide differential protein composition in neuronal cell types and thus help solidify each of the numerous cellular identities found in the brain. In the mouse hippocampus, sc-LR-Ribo-STAMP measurements revealed a possible role for retained introns in contributing to cell-type-specific gene expression regulation^15^. The same study found that the long 3′UTRs isoforms of individual synaptic genes, which were enriched in nELAV binding sites, translated more efficiently than their short counterpart. Since the expression of ELAV/Hu factors correlates with long UTRs in Drosophila^21,68,69^ and mouse^70^ brains, it is possible that ELAV/Hu-mediated nUTR synthesis coupled to differential nUTR translation contributes to the establishment and maintenance of distinct neuronal cell types. Increasing the depth and resolution of technologies that combine single-cell translational profiling with long-read sequencing will be useful in understanding how nUTR-mediated translational repression helps refining the functional neuronal proteome, eventually at the proteoform level. Other approaches such as spatial multiomics, capable of detecting distinct transcript variants and their product at near-single-cell resolution, could soon shed some light on how mRNA isoform diversity shapes neuronal cell type identity.

## Methods

### Fly husbandry and mutagenesis

All flies were maintained at 25°C. All experiments on adult fly heads used 3–5-day-old flies, and larval brains were from wandering third instar larvae.

#### Strains generated in this study

CRISPR-Cas9 genome editing followed the procedure described by Port and Bullock^71^. To generate the *elav^ΔnUTR^* mutant, two guide RNAs targeted the *elav* 3′UTR; the resulting cut was repaired using a 1,773-nt ultramer as a homology donor (gBlock from IDT), generating a 3 kb deletion. Flies denoted as Control are progeny of a non-mutated sibling of the parental *elav^ΔnUTR^* fly. To generate the *elav* “Proximal PAS” mutant, two guide RNAs targeted the *elav* 3′UTR, and used a 768-nt ultramer as a homology donor (Eurofins) to replace the canonical proximal PAS signal sequences (AATAAA) with non-functional variants (AACAAA). Two PAS sequences were mutated in this strain, located 2,341 and 2,549-nt downstream of the *elav* stop codon. To generate the *elav “Ribozyme all”* and *“Ribozyme long”* mutants, a single guide RNA was used in each case to target the short or long 3′UTR of *elav*, respectively; the resulting cut was repaired using a homology donor (500 and 538 nucleotides, respectively; gBlocks from IDT) carrying the hammerhead ribozyme sequence described by Tuck and Bühler^40^. All gRNA and homology donor sequences are listed in Table S5. Increase of *elav* gene dosage was performed using BAC transgenes. The 97 kb *elav* CHORI BAC CH321-35G14 was used as control transgene. Recombineering was performed on CH321-35G14 to construct the *elav^ΔnUTR^* BAC using the recombineering protocol described in Venken et al.^72^. Primers used for recombineering are listed in Table S5. BACs were integrated in the landing site VK33 on chromosome 3. All Drosophila embryo injections were performed by Bestgene, Inc.

#### Strains obtained from external sources

*w^1118^* control flies (RRID: BDSC_5905) and GFP-marked balancer lines (RRID: BDSC_4559, BDSC_6663) were obtained from the Bloomington Drosophila Stock Center. *Δelav* flies are described in Carrasco et al.^22^. *elav^FLAG^*flies, which express an endogenously, N-terminally Flag-HA-tagged ELAV protein, were described in Alfonso-Gonzalez and Shi et al.^73^. The *pumilio* lack-of-function alleles *pum^ET^*^7^ and *pum^ET^*^9^ were kindly provided by Ruth Lehmann^74^ and crossed to obtain trans-heterozygous flies (denoted as *Δpum*), which survived until the third instar larval stage.

### Human iPSC gene editing

CRISPR-Cas9 genome editing followed the procedure described in the Integrated DNA Technologies instruction manual. The human iPSC line SCTi-003A was obtained from STEMCELL Technologies (200-0511). To generate the *ELAVL1^ΔnUTR^* (c.*1365_*3419del) cell line, cells were detached at 70% confluency using TrypLE Express (Gibco, 12604013). 5×10^5^ cells were used per reaction with a NEPA21 Type2 electroporator (NepaGene, NEPA21). Alt-R™ S.p. Cas9 Nuclease V3 (IDT, 1081058) and two sgRNAs designed using the IDT Alt-R Custom Cas9 Design Tool were transfected in 1:3 ratio (sequences in Table S5). Electroporation was performed with the following settings: Poring Pulse 125V, Length 2.5 ms, Interval 50ms, Number 2, Decay rate 10%, Polarity +; Transfer Pulse 20V, Length 50 ms, Interval 50ms, Number 5, Decay rate 40%, Polarity +/-. Cells were transfected with RNP complexes and StemMACS™ eGFP mRNA (Miltenyi Biotec, 130-101-114). After two days, GFP-positive cells were single-cell sorted and screened for the genomic deletion.

### Neuronal induction of human iPSCs

Neuronal differentiation was performed using a protocol adapted from Chambers et al.^75^. Briefly, Human induced pluripotent stem cells (iPSCs) were cultured in mTeSR1 complete medium (STEMCELL Technologies, 100-0276) and seeded at 4×10^5^ cells/well on Matrigel-coated 12-well plates. Dual SMAD inhibition was initiated on day 1 by supplementing mTeSR1 with SB431542 (10 µM; Selleck Chemicals, S1067) and Dorsomorphin (1 µM; Reprocell, 04-0024). Thiazovivin (2 µM; STEMCELL Technologies, 72252), a ROCK inhibitor, was used to promote cell survival following single-cell dissociation. From day 6, the mTeSR1-based medium was gradually replaced with DMEM/F12 (Thermo Fisher Scientific, 31331093) supplemented with N2 (Gibco, 17502048) to promote growth while maintaining SMAD inhibition, transitioning fully to a DMEM-based medium by day 9. On day 13, cells were harvested and seeded at high density (1.4× 10^6^ cells/12-well) in hNSC growth medium (DMEM/F12, N2, 20 ng/mL bFGF (PeproTech, AF-100-15-500UG), 20 ng/mL EGF (PeproTech, 100-18C-100UG)) supplemented with 2% FBS (Gibco, A5256701) and 2 µM Thiazovivin for an explicit neural stem cell maintenance stage. On day 15, terminal neuronal differentiation was induced on Poly-L-Ornithine/Laminin coated plates using Neurobasal medium (Gibco, 21103049) supplemented with 1:50 B-27 and 1:100 GlutaMAX. All collections were performed in triplicate, with each replicate derived from a separate well of a 12-well plate. NSCs were collected 2 days after splitting into NSC medium (corresponding to day 15), and neurons were collected 15 days after neuron induction (day 30).

### Human cerebral organoid generation and processing

Human cerebral organoids (hCOs) were generated according to an adapted version of the protocol described by Lancaster et al.,^76^, as implemented by Gather et al.^77^. Briefly, 35,000 hiPSC cells per well, maintained in mTeSR Plus medium (STEMCELL Technologies, 100-0276), were centrifuged and seeded into V-shaped, non-coated 96-well plates and treated as described^77^. Organoids were cultivated for up to 34 days (d34), counted from the day of transfer from 96-well plates into petri dishes maintained on a horizontal shaker at 50 rpm. Half of the medium was changed once per week until d34. On d5, organoids were harvested and processed for lysate preparation for western blot analysis. On d34, organoids derived from control and *ELAVL1^ΔnUTR^* lines were processed for RNA extraction and cryosectioning for immunostaining analysis. For RT-qPCR and western blot, three organoids were pooled per biological replicate. To monitor organoid growth, images were captured with an Olympus CKX53 light microscope. Organoid cryogenic tissue processing and section immunofluorescence followed STEMCELL Technologies’ protocol, with the following specifications. Cryosections were dried (30 min, 50°C), washed in PBS, and subjected to antigen retrieval in sodium citrate buffer (10 mM, Tween-20 0.05%, pH 6.0) for 1 h at 60–70°C. Sections were quenched with 0.1 M glycine, blocked in 10% donkey serum with 0.01% Triton X-100, and incubated with primary antibodies diluted in PBS containing 0.01% Triton X-100 overnight at 4°C. Primary antibodies were anti-FOXG1 (Abcam, Ab18259, 1:100), rabbit anti-SOX2 (Cell Signaling Technology, 23064S, 1:200), rabbit anti-NeuN (Invitrogen PA-78499, 1:200), rabbit anti-TuJ1 (Proteintech, 10094-1-AP, 1:50), and mouse anti-ELAVL1 (Proteintech, 66549, 1:50). Sections were washed in PBS and incubated with secondary antibodies (Thermo Fisher Scientific, donkey anti-rabbit Alexa Fluor 488 and donkey anti-mouse Alexa Fluor 594, 1:500) for 1 h at room temperature, washed and counterstained with DAPI, and mounted using Fluoromount (Thermo Fisher Scientific, 00-4958-02).

### Lysate Preparation for polysome profiles

#### Fly heads

Flies were anesthetized and were blended using Philips Pro Blend 6 kitchen blender in cold PBS supplemented with 50 µg/mL cycloheximide (Thermo Fischer Scientific J66665.06). The homogenate was poured through pre-cooled sieves with aperture sizes 710 µm–425 µm (top to bottom) using pressurized cold distilled water. Fly heads were collected on the 425 µm sieve and transferred to a tube with 5 mL homogenization buffer (20 mM HEPES (pH 7.4), 120 mM potassium acetate, 2 mM magnesium acetate, 50 µg/mL cycloheximide, protease inhibitor cocktail (Roche, 11873580001), and RiboLock RNase inhibitor (Thermo Fisher Scientific, EO0384)), then homogenized using a Heidolph digital laboratory stirrer (RZR 2021, 400 rpm, 1 dounce). The homogenate was passed through a 40 µm strainer with the help of a sterile spatula and was centrifuged at 1000 g at 4°C to pellet down debris and nuclei. The supernatant was centrifuged at 12,000 g for 30 min at 4°C. The supernatant was subjected to a clearing spin for 1 min at 12,000 g in 4°C. 200 µL final lysate, containing 300–400 µg RNA, were loaded onto 11 mL linear sucrose gradient (15–55%, prepared in homogenization buffer using Biocomp gradient maker) by slowly dispensing it along the walls of the ultracentrifuge tubes, followed by ultracentrifugation at 200,000 g (TH641, Sorvall) for 2 h, and fractionated into 560 µL aliquots with a gradient fractionator monitoring A_260_ (Piston Gradient Fractionator, Biocomp). 50 µL of lysate (input) and each relevant polysomal fraction were dissolved in TRIzol LS Reagent (Ambion, 10296028) for RNA extraction.

#### Mouse brains

Whole brains from 8–13-week-old male and female C57BL/6J mice were dissected, washed thrice with ice-cold PBS, submerged in 2 mL Neurobasal™ medium (Gibco, 21103-049) containing cycloheximide (100 µg/mL), and incubated at 37°C for 10 min. Tissues were centrifuged at 500 g for 5 min at 4°C, the supernatant was discarded, and tissues were washed twice with ice-cold PBS containing 100 µg/mL cycloheximide. Tissues were resuspended in 500 µL of homogenization buffer (20 mM HEPES pH 7.4, 120 mM potassium acetate, 2 mM magnesium acetate, 50 µg/mL cycloheximide, 1% Triton X-100, 1 mM DTT, 0.50% sodium deoxycholate, protease inhibitor cocktail (Roche, 11873580001), RiboLock RNase inhibitor (Thermo Fisher Scientific, EO0384)) and gently lysed using an insulin needle. The homogenate was incubated on ice for 10 min with brief vortexing every 2–3 min and centrifuged at 2,000 g for 5 min at 4°C. The supernatant was transferred to a new tube on ice, and centrifuged again at 13,000 g for 5 min at 4°C. The clarified supernatant (final lysate) was transferred to a fresh tube on ice and all samples were diluted to roughly equal RNA concentrations with additional homogenization buffer. 250 µL of lysate (containing 400–500 µg total RNA) was carefully layered onto a 13 ml 20–60% linear sucrose gradient prepared in the homogenization buffer using a BioComp Instruments gradient maker. Gradients were subjected to ultracentrifugation at 218,000 g for 2.5 h at 4°C using an SW-40 Ti rotor (Beckman Coulter). Following centrifugation, gradients were fractionated into 150 µL aliquots using a Piston Gradient Fractionator (Gilson/BioComp Instruments), with continuous monitoring of absorbance at A_260_ and A_280_. 50 µL of lysate (input) and each relevant polysomal fraction were dissolved in TRIzol LS Reagent (Ambion, 10296028) for RNA extraction.

#### Human NSCs

hNSCs grown to 80–100% confluency in 15 cm dishes (approximately 20 million cells per dish) were harvested, with each dish constituting one biological replicate (three replicates in total). Prior to harvesting, cells were treated with 100 µg/mL cycloheximide for 5 min at 37°C to stabilize ribosome–mRNA complexes. Cells were harvested by scraping at 4°C and transferred into 15 mL conical tubes with 1.5 mL neurobasal medium (Gibco, 21103049) supplemented with 100 µg/mL cycloheximide. The cell suspension was centrifuged at 200 g for 5 min, the supernatant was discarded, and the pellet was resuspended in 500 µL homogenization buffer (20 mM HEPES (pH 7.4), 120 mM potassium acetate, 2 mM magnesium acetate, 50 µg/mL cycloheximide, 1% Triton X-100, 1 mM DTT, 0.50% sodium deoxycholate, protease inhibitor cocktail (Roche, 11873580001), RiboLock RNase inhibitor (Thermo Fisher Scientific, EO0384)). Lysates were centrifuged at 4,000 g for 10 min to remove nuclei and cell debris. The resulting supernatant was transferred to a fresh tube and subjected to an additional clearing spin at 12,000 g for 5 min. 250 µL of clarified supernatant (containing 400–500 µg total RNA) was carefully layered onto a 13 mL 20–60% linear sucrose gradient prepared in the homogenization buffer using a BioComp Instruments gradient maker. Gradients were subjected to ultracentrifugation at 218,000 g for 2.5 h at 4°C using an SW-40 Ti rotor (Beckman Coulter). Following centrifugation, gradients were fractionated into 150 µL aliquots using a Piston Gradient Fractionator (Gilson/BioComp Instruments), with continuous monitoring of absorbance at A_260_ and A_260_. 50 µL lysate (input) and each relevant polysomal fraction were dissolved in TRIzol LS Reagent (Ambion, 10296028) for RNA extraction.

### RNA extraction and RT-qPCR

Samples were dissolved in TRIzol LS (all polysome profiling fractions) or QIAzol Lysis Reagent (QIAGEN, 79306) and RNA was extracted according to the manufacturer’s instructions. Unless mentioned otherwise, 500 ng total RNA was used for cDNA synthesis using the iScript gDNA Clear cDNA Synthesis Kit (Bio-Rad). RT-qPCR was performed in a LightCycler 480 II instrument using FastStart SYBR Green Master Mix (Roche). RT-qPCR primer sequences are listed in Table S5.

### Western Blotting

For all western blots with adult Drosophila heads and larval brains, 10 heads per sample were homogenized in 50 or 100 µL sample buffer (1x NuPAGE (Invitrogen, NP0007) in PBS, 200 mM DTT), and 10 µL were loaded per lane. Primary antibodies rabbit anti-ELAV (homemade; described in^22^) and rabbit anti-histone H3 (Abcam, Ab1791; RRID:AB_302613) were used at concentrations 1:2,000 and 1:10,000 respectively. The secondary peroxidase-conjugated anti-rabbit antibody (Cell Signaling Technology, #7074; RRID:AB_2099233) was used at concentration 1:2,000. Band intensity for all western blot quantifications was measured using Fiji (ImageJ) or Image Lab 6.0.1 software.

### Starvation assay

The starvation protocol for flies was adapted from Service et al.^78^. Flies were maintained in vials either containing standard food (fed condition) or without food (starved condition) for 24 h unless otherwise indicated. In the starved condition, flies were provided with water-soaked Whatman filter paper to prevent dehydration. For western blot and RNA extraction, fly heads were collected from flies that were alive at indicated time point post-starvation. Each biological replicate consisted of 10 fly heads (five females and five males).

### Hatching, eclosion and viability assays

For all assays, 3–5-day-old flies were used for egg laying. The hatching rate was calculated as the fraction of embryos that successfully completed embryogenesis and hatched into first instar larvae. Dechorionated embryos coated with halocarbon oil were placed on agar plates in batches of 10 (n=10, e.g., 100 embryos per genotype). The number of hatched embryos was recorded every hour starting at 22-24h after egg laying (AEL) until 25-27h AEL, and one last time at 46–48h AEL. To assess developmental delays, females were allowed to lay eggs for 24h in vials. The parent flies were discarded and the embryos allowed to develop at 25°C. The number of eclosed flies was recorded every day at the same time from day 8 until day 11 AEL. To assess viability, virgin homozygous females of the *elav^ΔnUTR^*and control genotypes were crossed separately with wild-type (*w^1118^*) males and allowed to lay eggs for 24h in vials. The parent flies were discarded and the embryos allowed to develop for 12 days at 25°C. Adult viability was measured as the fraction of eclosed hemizygous males compared to the total number of flies eclosed from the respective crosses.

### RNA sequencing

For library preparation, for each sample, 100 ng of total RNA (fly head polysome experiment, Pum xRIP, control and *elav^ΔnUTR^* fly heads (in starvation and fed conditions)) or 150 ng RNA (mouse brain and NSCs polysome experiment) were used for library preparation with the Stranded Total RNA with Ribo-Zero Plus kit (Illumina) following the manufacturer’s instructions. Stranded RNA sequencing was performed on the NovaSeq 6000 platform (Illumina) with 2 × 100 bp paired-end reads. Sequencing data were processed using the RNA-seq module from snakePipes v2.4.3^79^, adding flags for --trim, -m “alignment-free,alignment.” Reads were mapped to the fly reference genome (Ensembl assembly release dm6), and the transcriptome reference annotation release 96^80^. For *Mus musculus*, reads were mapped to the mm10 reference genome and the transcriptome reference annotation release 97. For *Homo sapiens*, reads were mapped to the hg38 reference genome and the transcriptome reference annotation release 97. Quality control of RNA-seq reads was done using FASTQC^81^ (http://www.bioinformatics.babraham.ac.uk). To compare gene expression estimates across the different comparisons and samples, a variance-stabilizing transformation (VST) was applied using the DESeq2^82^ function vst() on raw gene count data from the different samples. The transformed data were used to compute a PCA using the DESeq2 function plotPCA() with standard parameters. All the heatmaps for 3′UTR coverage and changes were generated using deeptools 3.5.6 computeMatrix^83^ with the parameters “scale-regions -b 500 –skipZeros --binSize 25” or “scale-regions -b 500 --skipZeros --binSize 50” and then running the plotHeatmap function.

### Analysis of Pumilio binding to 3′UTRs

Pum xRIP was performed as described in Grzejda and Hess et al.^46^, and RNA-sequencing used total RNA-seq (not 3’-end sequencing) to allow for optimal 3′UTR quantification. 3′UTRs *in D. melanogaster* were counted in the Pum xRIP data with the parameters “-p -B --primary - Q 1 -s 2” in featureCounts^84^. Per gene, counts in the most proximal 3′UTR were classified as proximal counts, while all other 3′UTR counts were merged together as distal counts. The resulting counts were then used to perform a differential analysis between Pum xRIP and input, to measure Pumilio binding for each 3′UTR. 313 ELAV APA targets were compared against a random sample of 300 APA genes that are not ELAV targets. P-values were calculated with a Wilcoxon test, as the data is non-normally distributed.

### Gene ontology analysis

GO enrichment analysis was performed using the DAVID online server (version 2025). All individual groups were queried against the background of all genes detected for our analysis based on expression in respective lysate. GO terms with p-values <0.001 were classified as significant and visualized.

### Poly(A) site usage analysis

Distal PAS usage (dPAU) was calculated for polysome data in *D. melanogaster*, *M. musculus* and *H.* sapiens using the QAPA pipeline^36^, by first determining proximal PAS usage (pPAU), then computing dPAU as 1 - pPAU. QAPA calculates PAU scores (0-1) for all fractions (lysate, translating and non-translating). Lengthening was defined as pPAU in lysate/pPAU in fraction <0.4, shortening as pPAU in lysate/pPAU in fraction >1.6. Heatmaps of dPAU values across all polysome fractions, clustered by Ward’s method (clustering_method = “ward.D2”) were generated using the pheatmap package in R version 4.2.1.

### Gene set comparison under starvation stress

Stress-related gene sets, comprising all stress-responsive genes, protein folding genes, redox genes, and fatty acid metabolism genes, were retrieved from FlyBase using the indicated gene ontology terms. For each gene set and genotype, the expression log_2_FC between fed and starved conditions was compared. Only genes that were upregulated under starvation in both control and *elav^ΔnUTR^* genotypes were retained for analysis. Differences in log_2_FC between the two genotypes were assessed for statistical significance using a Wilcoxon signed-rank test.

### Proteomics sample preparation, data acquisition and processing

Fly heads (*elav^ΔnUTR^*) and larval brains (*Δpum*) were lysed in RIPA buffer supplemented with protease inhibitors. Cleared lysates (∼50 µg total protein) were reduced (5 mM TCEP; Sigma Aldrich, 51805-45-9), alkylated (40 mM 2-chloroacetamide), and treated with benzonase prior to SP3-based cleanup as described by Atinbayeva et al^85^. Proteolytic digestion was performed in 50 mM ammonium bicarbonate with LysC (100 ng, 2 h, 37°C) followed by trypsin (500 ng, overnight, 37°C). Peptides were acidified, partially dried in vacuo, resuspended in 0.1% formic acid, and loaded onto Evotips (Evosep) as per manufacturer’s instructions. Samples were measured on a Bruker timsTOF HT coupled to an Evosep One HPLC using an 88-minute gradient. *elav^ΔnUTR^*experiment samples were acquired using DIA-PASEF (ion mobility 1/K0 0.6–1.6 V·s/cm², m/z 400–1200, 16 ramps/cycle). *Δpum* experiment samples were acquired using DIA Thin-PASEF^86^ with a narrowed ion mobility window (1/K0 0.7–1.3 V·s/cm², m/z 400–1200, 9 ramps/cycle). Raw data were analyzed with DIA-NN (v2.1.0 for *elav^ΔnUTR^*; v2.3.2 for *Δpum*) against a UniProt *D. melanogaster* database including isoforms appended with a common contaminants list. Search parameters included up to one missed cleavage, variable methionine oxidation, peptide length 7–40 AA, and 15 ppm mass tolerance at MS1 and MS2. Downstream processing was performed in R using the in-house package ProteomeR. Contaminants were removed and protein entries were filtered by Q.Value ≤ 0.01, PG.Q.Value ≤ 0.05, Lib.Q.Value ≤ 0.01, Lib.PG.Q.Value ≤ 0.01, and PG.MaxLFQ.Quality ≤ 0.8 (*elav^ΔnUTR^* experiment) or ≤ 0.7 (*Δpum* experiment). Proteins were retained if they had valid values in at least n-1 replicates (*elav^ΔnUTR^* experiment) or 60% of replicates (*Δpum* experiment) in any condition. Missing values were imputed as follows: proteins absent or missing in all replicates of a condition or in more than 60% (*elav^ΔnUTR^*experiment) or 75% (*Δpum* experiment) of samples were imputed by left-censored imputation (scale = 0.25, shift = 2.1 *elav^ΔnUTR^* experiment) or 1.9 (*Δpum* experiment)^87^). Remaining missing values were imputed using missForest^88^. Differential abundance was assessed using limma with empirical Bayes moderation (TREND = TRUE)^89^. For the *Δpum* dataset, one control replicate exhibiting a batch effect was included as a covariate in the linear model. P-values were Benjamini-Hochberg adjusted. Proteins were considered differentially abundant at adjusted p ≤ 0.05 and fold change ≥ ±50% (*elav^ΔnUTR^*) or ≥ ±25% (*Δpum*).

## Supporting information

Table S1 part 1

Table S1 part 2

Table S1 part 3

## Data availability

All information and requests for resources and reagents used and/or generated in this study are available on reasonable request from the corresponding author with a completed materials transfer agreement. All sequencing data generated during this study (total RNA-seq data of polysome fractionation experiment in fly heads, mouse brains and NSCs; total RNA-seq in fly heads post starvation stress (control and *elav^ΔnUTR^*); and Pum xRIP-seq) can be accessed at NCBI Gene Expression Omnibus under the accession number GSE320358. Mass spectrometry data have been deposited to ProteomeXchange via PRIDE (*elav^ΔnUTR^*: PXD077804, *Δpum*: PXD077875).

## Acknowledgements

We thank Ulrike Bönisch at the MPI-IE Deep Sequencing Core, Gerhard Mittler at the MPI-IE Proteomics Core, and the MPI-IE Bioinformatics Core. We thank Ritwick Sawarkar, Giorgos Pyrowolakis, and Cong Truc Le for helpful discussions and Baivabi Bhattacharya, Gunashree Rathi, Evrim Selen Demirbasoglu, and Begoña Nieto Fernández for technical assistance. Stocks obtained from the Bloomington Drosophila Stock Center (NIH P40OD018537) were used in this study. J.C. is currently employed at Astrazeneca UK Ltd. This work was funded by the Max Planck Society, the German Research Foundation (DFG) SFB 1381 (Project-ID 403222702), the European Research Council (ERC) under the European Union’s Horizon 2020 research and innovation programme (ERC-2018-STG-803258), and ERC-2024-COG-NeuroRNA funded by the Swiss State Secretariat for Education, Research and Innovation (SERI).

## Contributions

V.H., S.G., J.C. Investigation and Validation: S.G., J.C., Y.Z., A.H., I.A., D.G., F.M., S.H., V.H. Formal analysis: S.G., H.C.O., A.H., C.A.-G., M.S., A.G.A. Visualization: S.G., H.C.O, C.A.-G., V.H. Methodology: S.G., Y.Z., I.A., J.M., M.E., A.G.A. Writing—original draft: S.G., V.H. Writing—review and editing: S.G., V.H., H.C.O., A.H. Supervision: V.H., N.C.-W., T.V., S.R. Funding acquisition: V.H.

## Ethics declarations

The authors declare no competing interests.

