## Supplementary Figures for "Regulation of protein abundance in neurons by selective translation of 3′UTR isoforms"

#### Supplemental Tables

**Table S1. QAPA analysis: differential translation of short and long 3'UTR isoforms for each gene in fly heads, mouse brains, and human NSCs.**

**Table S2. Gene Ontology analysis for genes with differential translation short and long 3'UTR isoforms in Drosophila heads and mouse brains.** As background gene set, all genes expressed in the respective tissue lysate were used.

**Table S3. Mass spectrometry analysis of protein expression in control and *elav*$^{\Delta nUTR}$ fly brains.**

**Table S4. Mass spectrometry analysis of protein expression in brains of control and *Δpum* third-instar ("wandering") larvae.**

**Table S5. Nucleotide sequences used in this study.**

### Supplemental Figures

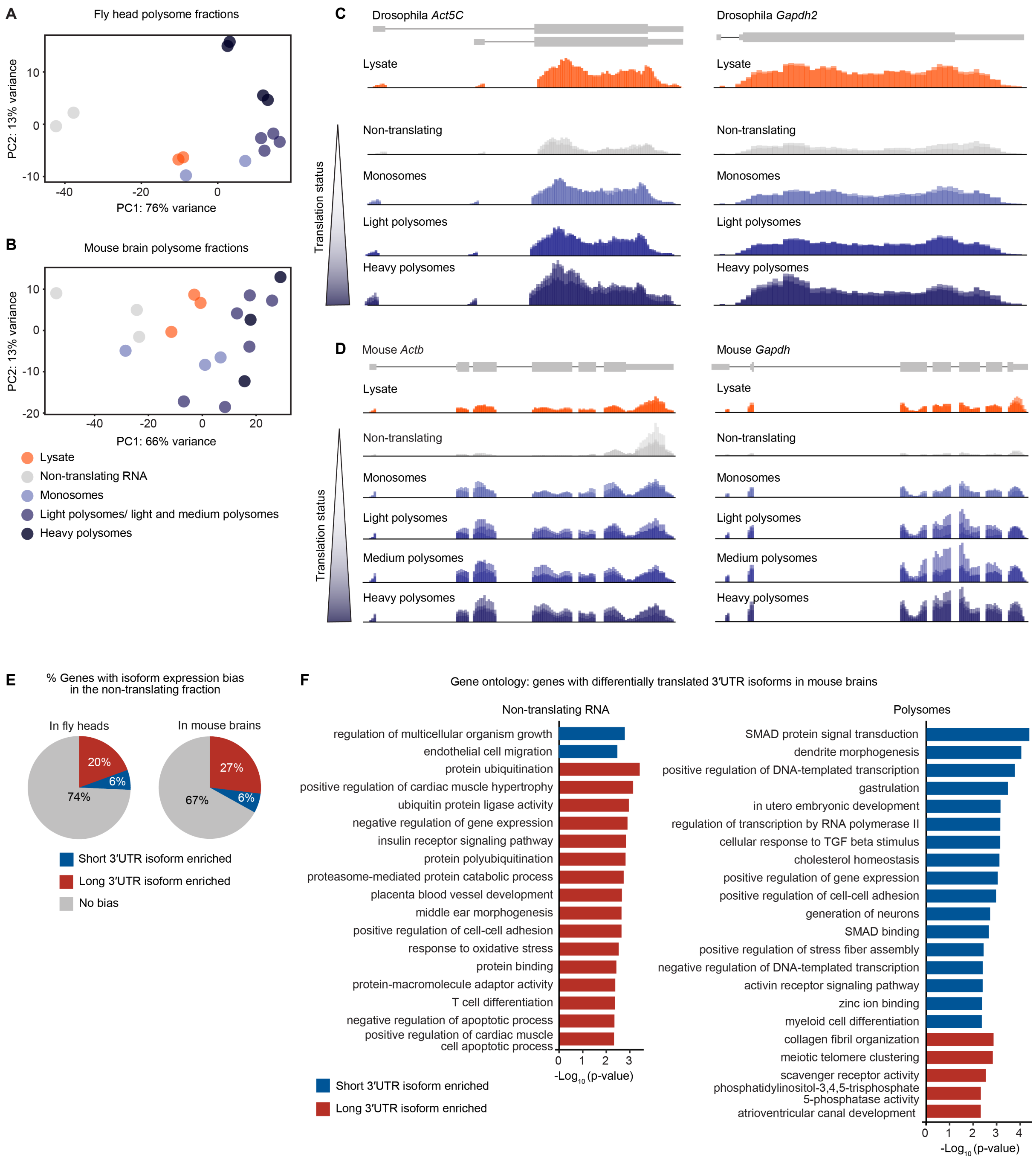


**Figure S1. Long 3′UTR isoforms of neuronal genes are less translated than their short counterparts.**

**(A, B)** Principal component analysis of gene expression in lysate and polysome profile fractions from fly heads **(A)** and mouse brains **(B)**.

**(C, D)** Total RNA-seq tracks of the indicated genes in lysate and polysome profile fractions from fly heads **(C)** and mouse brains **(D)**.

**(E)** Distribution of short, long 3'UTR or no isoform expression bias in the non-translating polysome fraction as compared to lysate, in all expressed genes with more than one polyadenylation site.

(**F**) Gene ontology analysis of genes for which short (blue) or long (red) 3'UTR isoforms are enriched in either translating or non-translating fractions from mouse brains. Background set: all expressed genes.

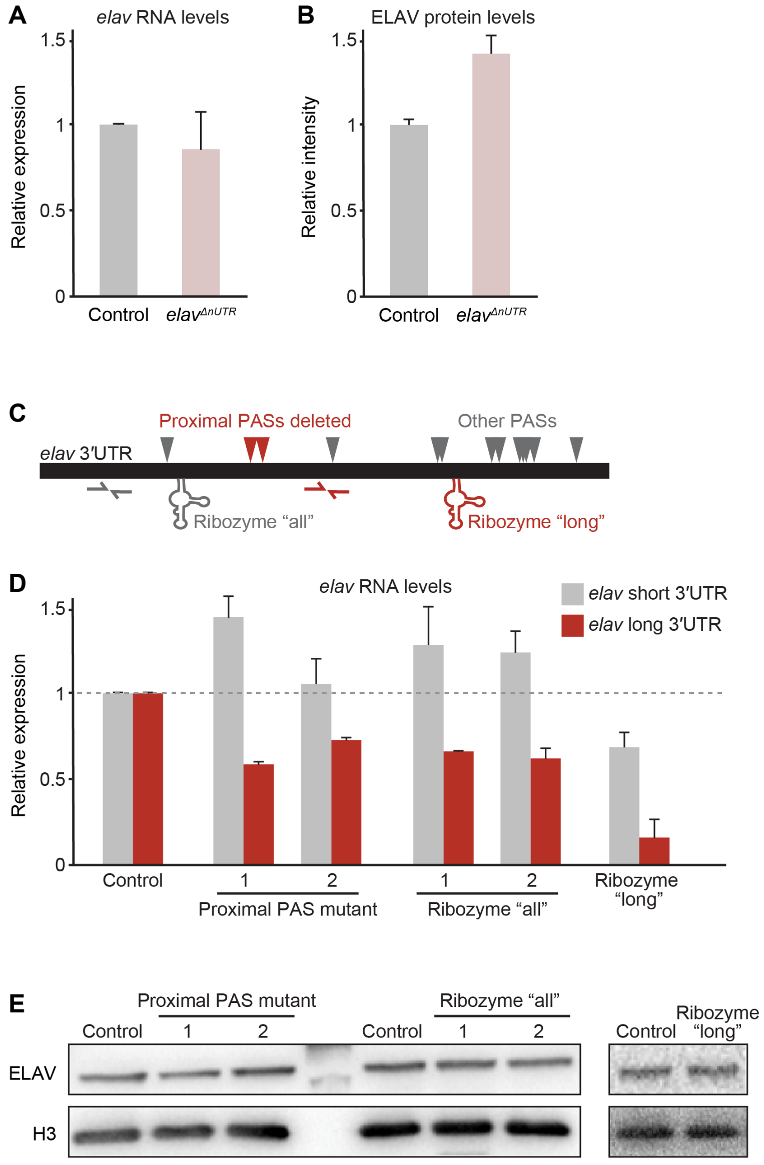


**Figure S2. The neuronal 3'UTR regulates mRNA translation and maintains optimal ELAV protein levels.**

**(A)** RT-qPCR quantification of *elav* mRNA levels in control and $elav^{\Delta nUTR}$ adult fly heads. RNA levels were normalized to *RpL32* mRNA and levels in control flies were set to 1. Error bars represent mean ±SD of three biological replicates for each genotype.

**(B)** Quantification of ELAV protein expression in control and $elav^{\Delta nUTR}$ adult fly heads as measured by mass spectrometry. Error bars represent mean ±SD of three biological replicates for each genotype.

**(C)** Schematic of the *elav* 3'UTR sequence, with all polyadenylation sites (PASs) indicated with arrowheads. Three distinct mutations generated through gene editing are represented. In "proximal PAS" mutant flies, canonical poly(A) signals of the two PASs indicated in red were mutated (AATAAA changed to AACAAA). In "Ribozyme all" and "Ribozyme long" flies, self-cleaving hammerhead ribozyme sequences were inserted proximally or distally, respectively, of the proximal PASs in the endogenous *elav* gene locus, to cause the degradation of all ("all") or specifically the long 3'UTR isoforms ("long") upon transcription of the ribozyme sequence. Half-arrows represent primer pairs used in the RT-qPCR shown in **(D)**.

**(D)** RT-qPCR quantification of the short (grey) and long (red) *elav* 3'UTR isoforms in adult fly heads of the indicated genotypes. RNA levels were normalized to *RpL32* mRNA and levels in control flies were set to 1. Error bars represent mean ±SD of three biological replicates for each genotype. Control flies are progeny of a non-mutated sibling of the parental mutagenized fly.

**(E)** ELAV protein expression in adult fly heads of the indicated genotypes. Histone H3 serves as a loading control. Two independent mutants are shown for the genotypes "proximal PAS mutant" and "Ribozyme all".

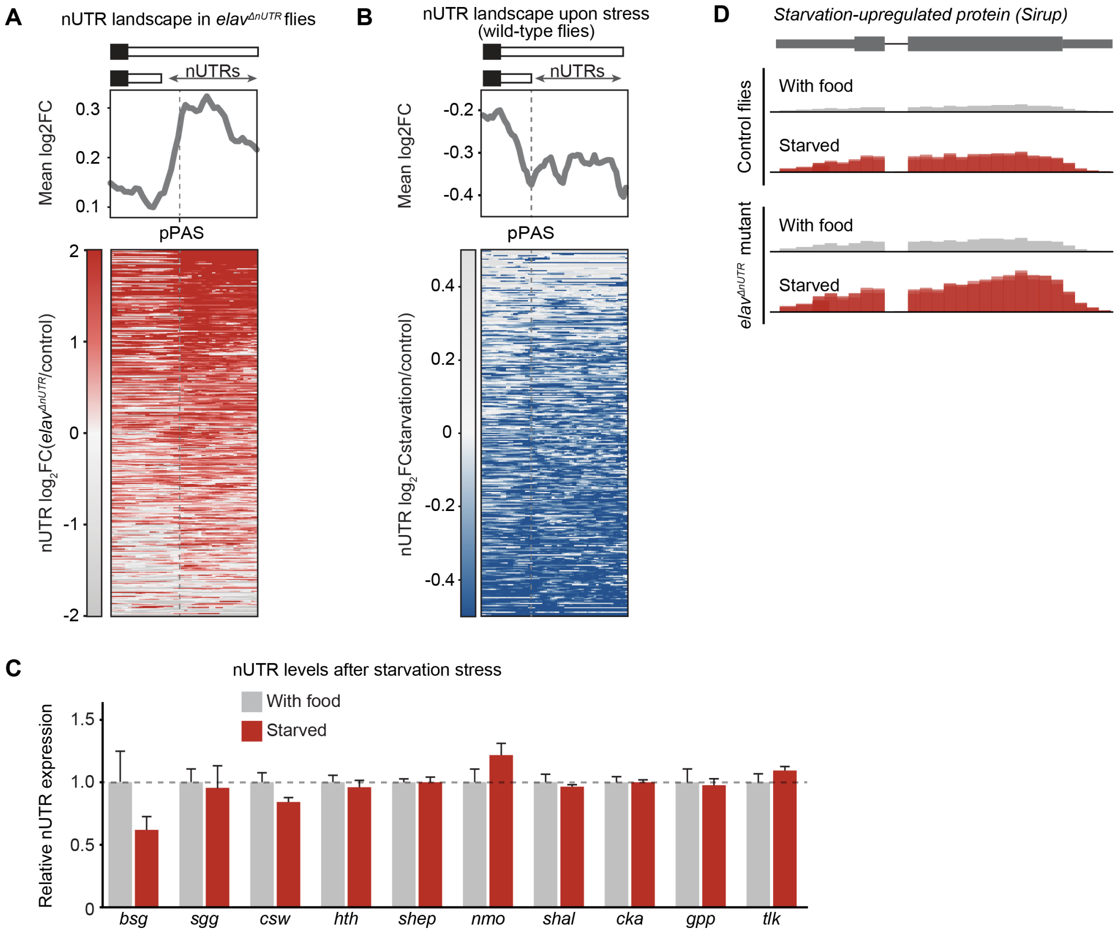


**Figure S3. 3′UTR-dependent translational regulation maintains proteostasis during development and stress.**

**(A)** Enrichment of long, neuron-specific 3′UTRs (nUTRs) in *elav*$^{\Delta nUTR}$ adult fly heads compared to control fly heads. The heatmap and profile plot display 0.5 kb upstream, and a portion of the nUTR downstream (scaled region), of the proximal poly(A) site.

**(B)** Differential nUTR expression in heads of adult flies kept under starvation stress for 24h compared to control fly heads. The heatmap and profile plot display 0.5 kb upstream, and a portion of the nUTR downstream (scaled region), of the proximal poly(A) site.

**(C)** RT-qPCR quantification of the nUTR-containing mRNA isoform relative to total (short) mRNA isoforms of the indicated genes, in adult heads of flies kept in control (with food) conditions or under starvation stress for 24h (starved). nUTR levels were normalized to respective coding sequence levels for each gene and levels in control conditions were set to 1. Error bars represent mean ±SD of three biological replicates.

**(D)** Total RNA-seq tracks of the gene *up on starvation* (*sirup*) in adult fly heads in the indicated genotypes and conditions.

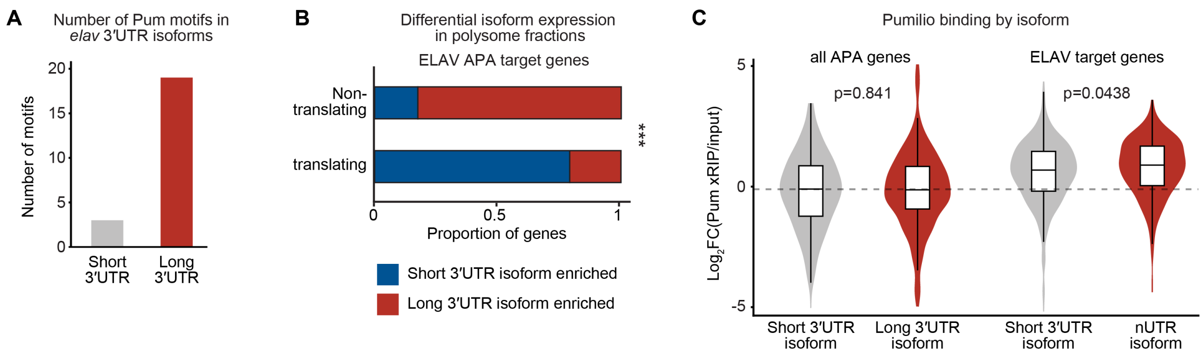


**Figure S4. Mechanism of nUTR-dependent translational repression.**

**(A)** Number of Pum motifs (TGTANWT) in the 3'UTR of the *elav* short and long 3'UTR isoforms.

**(B)** Fraction of genes with a 3′UTR isoform expression bias towards the short or long 3′UTR in the indicated polysome fractions compared to lysate. All genes expressed in adult fly brains that undergo ELAV-mediated 3'UTR extension (ELAV APA target genes) were considered for this analysis.

**(C)** Quantification of Pum binding to the long and short mRNA isoforms of the indicated gene groups, by Pum xRIP-seq signal compared to input. Data for ELAV target genes are from Fig. 4 and reproduced here for comparison. P-values by Wilcoxon test.

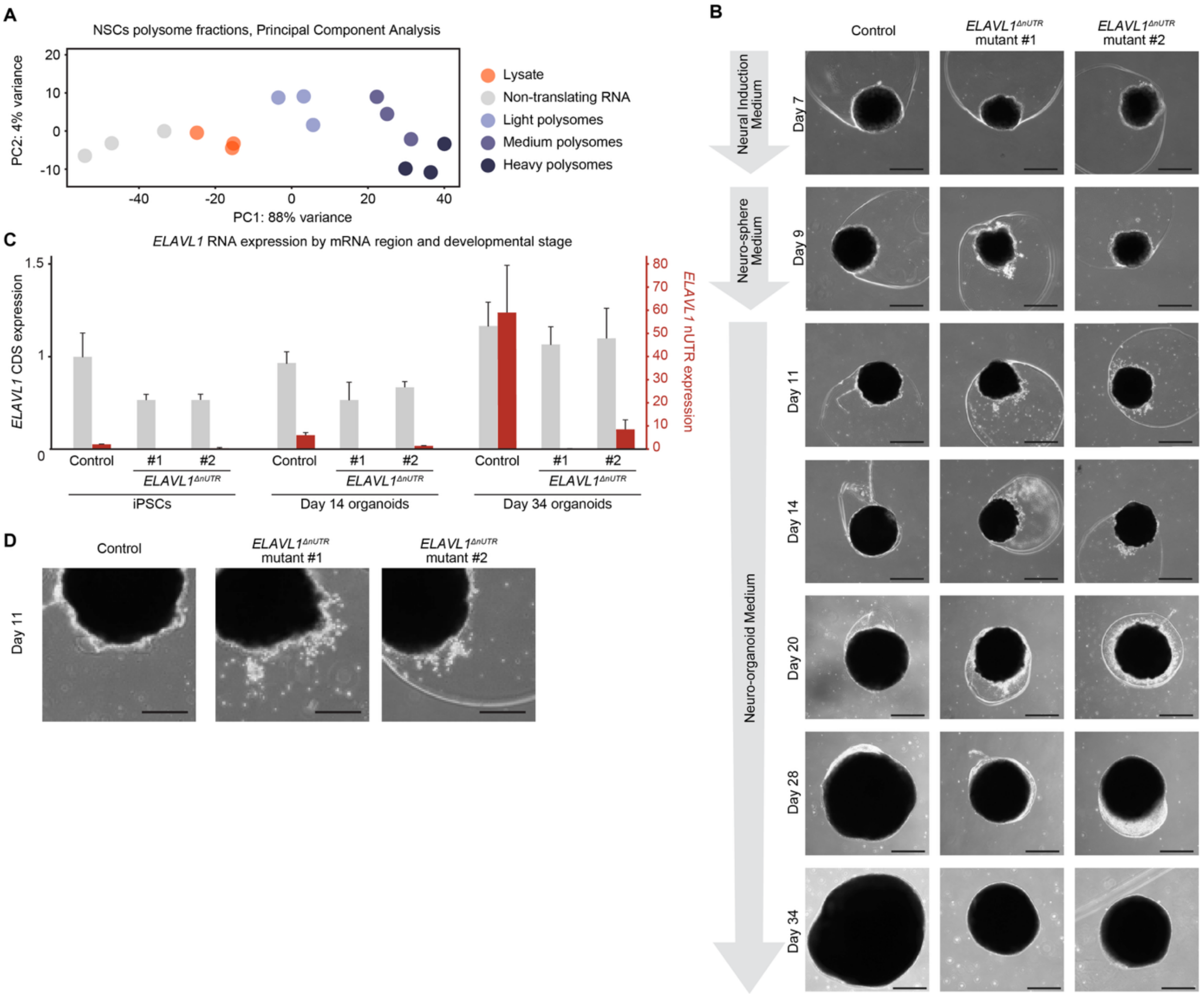


**Figure S5. Neurodevelopmental impairment in human brain organoids lacking the *ELAVL1* nUTR.**

**(A)** Principal component analysis of gene expression in lysate and polysome profile fractions from NSCs.

**(B)** Light microscopy images of cerebral organoids grown from control and *ELAVL1*$^{\Delta nUTR}$ human iPSCs, at the indicated number of days post-seeding. Scale bar: 500 µm.

**(C)** RT-qPCR quantification of *ELAVL1* coding sequence (left Y-axis, grey) and nUTR (right Y-axis, red) in control and *ELAVL1*$^{\Delta nUTR}$ (two independent mutants) iPSCs, day 14 organoids, and day 34 organoids. RNA levels were normalized to *GAPDH* mRNA and levels in control iPSCs were set to 1. Error bars represent mean ±SD of three biological replicates.

**(D)** Magnified image cutouts of control and *ELAVL1*$^{\Delta nUTR}$ day 11 organoids. Scale bar: 250 µm.
